# Major role of S-glycoprotein in providing immunogenicity and protective immunity in mRNA lipid nanoparticle vaccines based on SARS-CoV-2 structural proteins

**DOI:** 10.1101/2023.12.30.573713

**Authors:** Evgeniia N. Bykonia, Denis A. Kleymenov, Vladimir A. Gushchin, Andrey E. Sinyavin, Elena P. Mazunina, Nadezhda A. Kuznetsova, Sofia R. Kozlova, Anastasia N. Zolotar, Elena V Shidlovskaya, Evgeny V. Usachev, Andrei A. Pochtovyi, Daria D Kustova, Igor A. Ivanov, Sergey E. Dmitriev, Roman A. Ivanov, Denis Y. Logunov, Alexander L. Gintsburg

**Affiliations:** National Research Centre for Epidemiology and Microbiology named after Honorary Academician N F Gamaleya of the Ministry of Health of the Russian Federation, Moscow, Russia; Department of Virology, Lomonosov Moscow State University, Moscow, Russia; Shemyakin-Ovchinnikov Institute of Bioorganic Chemistry of the Russian Academy of Sciences, Moscow, Russia; Belozersky Institute of Physico-Chemical Biology, Lomonosov Moscow State University, Moscow, Russia; Faculty of Bioengineering and Bioinformatics, Lomonosov Moscow State University, Moscow, Russia; Sirius University of Science and Technology, Sochi, Russian Federation; Infectiology Department, I. M. Sechenov First Moscow State Medical University, Moscow, Russia

## Abstract

Recently we have developed an mRNA lipid nanoparticle (mRNA-LNP) platform providing efficient long-term expression of an encoded gene *in vivo* after both intramuscular and intravenous application. Based on this platform, we have generated mRNA-LNP coding SARS-CoV-2 structural proteins M, N, S from different virus variants and studied their immunogenicity separately or in combinations *in vivo*. As a result, all candidate vaccine compositions coding S and N proteins induced excellent anti-RBD and N titers of binding antibodies. T cell responses mainly represented specific CD4+ T cell lymphocyte producing IL-2 and TNF-α. mRNA-LNP coding M protein did not show high immunogenicity. High neutralizing activity was detected in sera of mice vaccinated with mRNA-LNP coding S protein (alone or in combinations) against closely related strains but was not detectable or significantly lower against an evolutionarily distant variant. Our data showed that the addition of mRNAs encoding S and M antigens to the mRNA-N in the vaccine composition enhanced immunogenicity of mRNA-N inducing more robust immune response to the N protein. Based on our results, we suggested that the S protein plays a key role in enhancement of immune response to the N protein in the mRNA-LNP vaccine.

## INTRODUCTION

Since the beginning of the COVID-19 pandemic, there have been several global waves of increased incidence caused by the rapidly evolving SARS-CoV-2 variants: Alpha (B.1.1.7), Beta (B.1.351), Gamma (P.1), Delta (B.1.617.2), Omicron (B.1.1.529) and others. Mutations in the gene encoding the SARS-CoV-2 spike protein have been found to contribute to immune evasion and/or increased transmissibility of the virus, resulting in reduced effectiveness of existing approved vaccines that primarily target the viral S protein to produce a potent neutralizing antibody response [1][2][3].

WHO Technical Advisory Group on COVID-19 Vaccine Composition in WHO Global COVID-19 Vaccination Strategy, published in June 2022, called for future boosters to be compatible with circulating SARS-CoV-2 variants and/or have broad immunogenicity against new variants of concern [4]. As a result, in the fall of 2022 Pfizer/BioNTech and Moderna produced booster bivalent mRNA vaccines coding spike proteins of the original SARS-CoV-2 variant and one of the Omicron variants BA.1 or BA.4/5 [5][6]. These updates in vaccine antigen composition led to enhanced vaccine-induced immune responses to circulating SARS-CoV-2 variants [7].

However, soon after Omicron sublineage XBB was first identified in August 2022, the XBB.1 descendants (i.e., XBB.1.5, XBB.1.16, XBB.1.9) became dominant globally in the first half of 2023 [8][9][10].

As a result, sera from individuals who have received two, three, or four doses of index virus-based vaccines or a booster dose of a bivalent mRNA vaccine (containing BA.1 or BA.4/5) show substantially lower neutralizing antibody titers against XBB.1 descendent lineages, as compared to titers specific to the antigens included in the vaccine [10]. Consequently, new updated COVID-19 vaccines developed by Pfizer-BioNTech and Moderna were approved by the Food and Drug Administration (FDA) in mid-September 2023 and recommended by the CDC. These updated vaccines consist of a monovalent composition, with mRNA coding for the S protein of Omicron XBB.1.5 only [11].

However, there is ongoing and considerable genetic and antigenic evolution of SARS-CoV-2, posing new challenges for the global community. To date, new variants of the virus have already predominated, and highly mutated ones (i.e. BA.2.86) have also been isolated [12][13]. It is too early to tell whether the newly adapted boosters will protect against these variants. Preliminary data suggests the potential utility of the monovalent XBB.1.5 mRNA boosters, but this is an area of ongoing research [13].

Therefore, continually updating vaccines provides high levels of protection against severe disease and mortality caused by all circulating variants of SARS-CoV-2, but it declines over time due to ongoing evolution. One possible approach to improve the durability and breadth of protection is including more conserved antigens in the vaccine in addition to the S protein.

Natural infections are known to provide superior immunity and broad protection compared to vaccines [14]. T cell response to SARS-CoV-2 indicate that there are many potential CD4+ and CD8+ T cells specific for both structural and non-structural proteins of the virus. In particular, the M, S, and N proteins are codominant in natural infection, each being recognized by CD4+ and CD8+ T cells in the vast majority of human COVID-19 survivors [15][16][17]. T cell responses develop early and play a central role in the control of SARS-CoV- 2, correlating with protection [18]. Thus, the inclusion of more conserved antigens (e.g., N and M proteins) in the vaccine may provide broader protection and long-last immunity [19][20][21].

Recently we have developed an mRNA-LNP platform providing efficient long-term expression of an encoded gene *in vivo* after both intramuscular and intravenous application. The platform has been extensively characterized using firefly luciferase (Fluc) as a reporter [22]. Based on this platform, in this study we generated mRNA-LNP vaccine-candidates that encoded antigens (M, N, and S proteins) from different SARS-CoV-2 variants. We used gene sequences of two early variants (Wuhan and Delta) and two later ones (Omicron BA.5.3.1 and XBB.1). The aim of this study was to evaluate the immunogenicity and protective efficacy of each antigen encoded in mRNA vaccine alone or in combination with each other and make possible recommendations regarding vaccine composition against SARS-CoV-2 VOCs.

## RESULTS

In this study, we investigated immune responses caused by SARS-CoV-2 structural proteins (S, M, and N) encoded in mRNA-LNPs vaccine-candidates, separately or in different combinations, to explore whether they influence each other and to what extent they mediate immunity. In addition, spike proteins were derived from evolutionarily distant SARS-CoV-2 variants (two from earlier ones: the wild-type strain Wuhan and Delta, and two from later ones:

Omicron BA.5.3.1 and XBB.1) to study their possible different abilities to induce an immune response.

Based on our recently developed mRNA platform [22], we generated mRNA-LNP vaccines encoding the following full-length structural proteins derived from different SARS-CoV-2 variants:

- Spike (S) protein of wild-type strain (Wuhan),
- Spike (S) protein of Delta variant,
- Spike (S) protein of Omicron variant (BA.5.3.1),
- Spike (S) protein of Omicron variant (XBB.1),
- N protein of Omicron variant (BA.5.3.1),
- M protein of Omicron variant (BA.5.3.1).

The coding regions for investigated antigens were cloned into the linear bacterial plasmid pJAZZ-OK and supplemented with all the elements needed for the production of effectively translated mRNA molecules [22]. The transcribed mRNAs had following structures providing its stability and a high translation rate: the cap-1 structure at the 5′ end; a 100-nt long poly(A)-tail at the 3′ end; the 5′ and 3′ UTRs from the human hemoglobin alpha subunit (*HBA1*) mRNA; coding sequences (CDS) of protein of interest **(Fig. 1A)**. In addition, 100% uridines (U) were replaced with N1-methylpseudouridines (m^1^Ψ) to reduce mRNA immunogenicity. Thus, mRNAs encoding the full-length SARS-CoV-2 structural proteins were obtained **(Fig. S1A)**.

**Figure 1.**
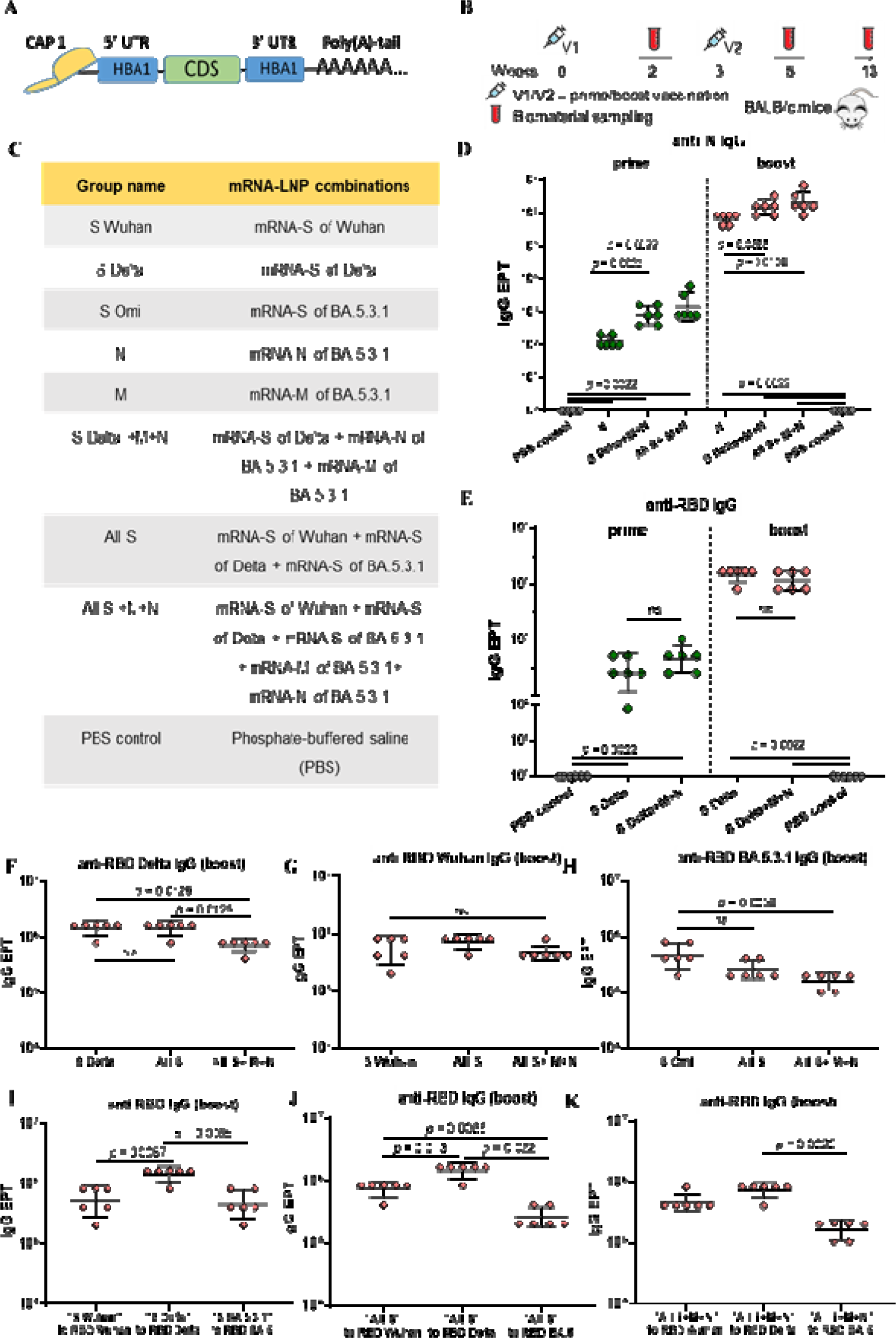
Structure of mRNA, composition of candidate vaccine compositions and assessment of humoral immunity. (**A**) Schematic representation of the mRNA platform used in this study. (**B**) Experimental design of immunogenicity assessment. Nine groups of BALB/c mice (*n* = 7 per group) were intramuscularly vaccinated with mRNA-LNP combination (5 μg of each mRNA per dose) or PBS (control group) at weeks 0 and 3. Two weeks after prime (V1) and booster (V2) vaccinations, serum samples were collected for analysis of binding antibody responses. Two and a half months after the booster mice were euthanized, serum and spleens were collected to determine virus neutralizing activity and T cell responses. (**C**) List of vaccine compositions and mouse group names. (**D**) Comparison of N-specific IgG end point titers (EPT) between PBS control group and vaccine groups after prime and booster vaccinations. (**E**) Comparison of RBD-specific IgG end point titers (EPT) between PBS control group and vaccine groups after prime and booster vaccinations. (**F-K**) Comparison of RBD-specific IgG end point titers (EPT) between vaccine groups after booster vaccination. EPT values are represented as scatter dot plots in logarithmic scale. Lines represents geometric mean with 95% confidence interval. Kruskal-Wallis test and the post hoc Dunn’s multiple comparisons test (F to H) or Mann-Whitney (D, E, I-K) test were used for statistical analysis (ns, not significant).

The protein expression ability of all mRNAs transcripts were examined in HEK293T cells. Cells were treated with mRNAs and then lysed. An in-house bead based immunoassay (Luminex® xMAP® technology) was used to detect desired proteins. Corresponding monoclonal antibodies (specific to N, M proteins and RBD domain of S protein) were conjugated to the magnetic beads according to the Cookbook [23]. The assay was performed in the form of a “sandwich” non-competitive immunoassay. Median fluorescent intensity (MFI) was measured on a MAGPIX instrument and converted in ng/ml. Results confirmed the expression of investigating proteins in cell culture, demonstrating the functionality of our mRNAs transcripts (**fig. S1B**).

Next step was formulating mRNAs in lipid nanoparticles by the microfluidic mixing procedure. Lipid mixture consisted of ionizable lipid (ALC-0315), DSPC, cholesterol, and PEGylated lipid (ALC-0159) at molar ratios of 46.3:9:42.7:1.6 respectively. Formulated mRNA-LNPs were characterized by particle size, polydispersity, zeta potential and mRNA encapsulation efficiency (**Table S1**). All mRNA-LNP sizes were below 100 nm (mean values 76, 75, 69 nm for mRNA-LNP coding S, N, M, respectively) with polydispersity index not exceeding 0.15 and zeta potential slightly negative but not less than ™10 mV. mRNA encapsulation efficiency averaged 86-91 %.

The immunogenicity of mRNA-LNP was evaluated in BALB/c mice. **Figure 1B** represents the scheme of the experiment. Eight groups of mice (n = 7 per group) were vaccinated with various combinations of mRNA-LNPs (5 μg of each mRNA-LNP per dose), phosphate-buffered saline was injected to the mice control group (PBS control). The mRNA-LNPs combinations for immunization and group names are listed in **figure 1C**. Immunization was conducted intramuscularly at week 0 (V1) and week 3 (V2). Two weeks after prime vaccination (V1) and two weeks after the booster (V2), serum samples were collected for analysis of binding antibody responses. Two and a half months after booster mice were euthanized, serum and spleens were collected to determine virus neutralizing activity and T cell responses (**fig. 1B**).

We determined binding IgG antibodies in the mouse serum samples after prime and boost immunizations by in-house enzyme-linked immunosorbent assay (ELISA). Antibodies specific to N, M proteins and RBD domain of S protein were analyzed (**fig.1 D-K**). For that, ELISA plates were coated by recombinant N or M proteins (1 μg/mL), or RBD-domains from investigating SARS-CoV-2 variants (Wuhan, Delta, BA.5.3.1) in the same concentrations (0.5 μg/mL). Antibody end point titers (EPTs) were determined in serially diluted serum samples.

Compared to the PBS control group, strong N- or/and RBD-specific binding IgG responses (*p* < 0.005) were observed in all groups immunized with mRNA-LNP coding N or/and S proteins, respectively, both after prime and boost vaccinations (**fig. 1D, E)**. Geometric mean titers (GMTs) of N-and RBD-specific IgG after boost dose were three orders of magnitude greater than after prime immunization in all mRNA-LNP vaccinated groups (*p* = 0.0022; *p* = 0.0048 (**fig. S2A**)) and reached maximum values: 1 459 648 for anti-RBD Delta IgG in groups “All S” and “S Delta” and 2 064 255 for anti-N IgG in “All S+M+N” group.

GMT for N- specific binding IgG in the group “N” was less than those for groups “S Delta+M+N” and “All S+M+N” both after prime (*p* = 0.0022) and boost (*p* < 0.005) doses (**fig. 1D**). The data suggest that the addition mRNAs encoding S and M antigens to the mRNA-N in the vaccine composition enhance immunogenicity of mRNA-N inducing more robust humoral response to N protein. This can be observed after both prime and boost vaccinations (**fig. 1D**). Conversely, comparing GMTs for RBD-specific binding IgG in the groups “S Delta” and “S Delta+M+N” did not reveal significant difference (**fig. 1E**). Thus, the addition of mRNAs encoding N and M proteins to mRNA-S vaccine preparation did not affect humoral response caused by mRNA-S (**fig. 1E**). Similarly, comparing GMTs for RBD-specific binding IgG in the groups “S Delta” and “All S” did not reveal significant difference (**fig. 1F**). Generally, combining of three mRNAs coding S protein of all used variant (Wuhan, Delta, Omicron BA.5.3.1) did not lead to an increase GMT of RBD-binding antibodies in “All S” group compared with those induced by alone mRNA-S (**fig. 1F, G, H**). However, vaccine composition comprising all used mRNAs (coding three S protein, N and M) contributed to lower GMT of RBD-binding IgG obtained for “All S+M+N” compared with those induced by alone mRNA-S of Delta (*p* = 0.0126) and mRNA-S of Omicron (*p* = 0.0058), but not mRNA-S of Wuhan (**fig. 1F, G, H**).

Furthermore, it was found that GMT of RBD-binding IgG induced in “S Delta” group was higher than those produced in “S Wuhan” (*p* = 0.0087) or “S Omi” group (*p* = 0.0065) (**fig. 1I**). In “All S” group, GMT of binding IgG specific to RBD of Delta was higher than GMT of binding IgG specific to RBD Wuhan (*p* = 0.013) or RBD Omicron BA.5.3.1 (*p* = 0.022) (**fig. 1J**). Similarly, in group “All S+M+N” GMT of binding IgG specific to RBD of Delta was higher than GMT of binding IgG specific to RBD Omicron BA.5.3.1 (*p* = 0.0006) (**fig. 1K**).

As for M- specific antibodies, poor responses were observed after both prime and boost immunizations in all mRNA-M vaccinated groups by the bead based immunologic analysis (ELISA analysis did not reveal any responses). M-specific binding antibodies were determined only in “M” group after booster compared with PBS control group (**fig. S3**).

Next, we evaluated virus-neutralizing activity in serum samples by the cytopathic effect (CPE) assay (**fig. 2**). High neutralizing activity was detected in groups vaccinated with alone mRNA-S (“S Wuhan”, “S Delta” and “S Omi” groups) against closely related strains (Wuhan-like strain B.1.1, Delta or Omicron BF.5, respectively) (**fig. 2A-D)**. GMT of neutralizing antibodies in these groups were 1 437, 1 613 and 735 respectively. However, neutralizing activity was not detectable (**fig. 2A, B)** or significantly lower (**fig. 2C**, *p* = 0.0079) against evolutionarily distant variant Omicron XBB.1.16. Similar results are demonstrated in groups “All S” and “All S+M+N” – high GMT values against B.1.1 and reduced activity for XBB.1.16 (**fig. 2D, H, G**, *p* = 0.0022)

**Figure 2.**
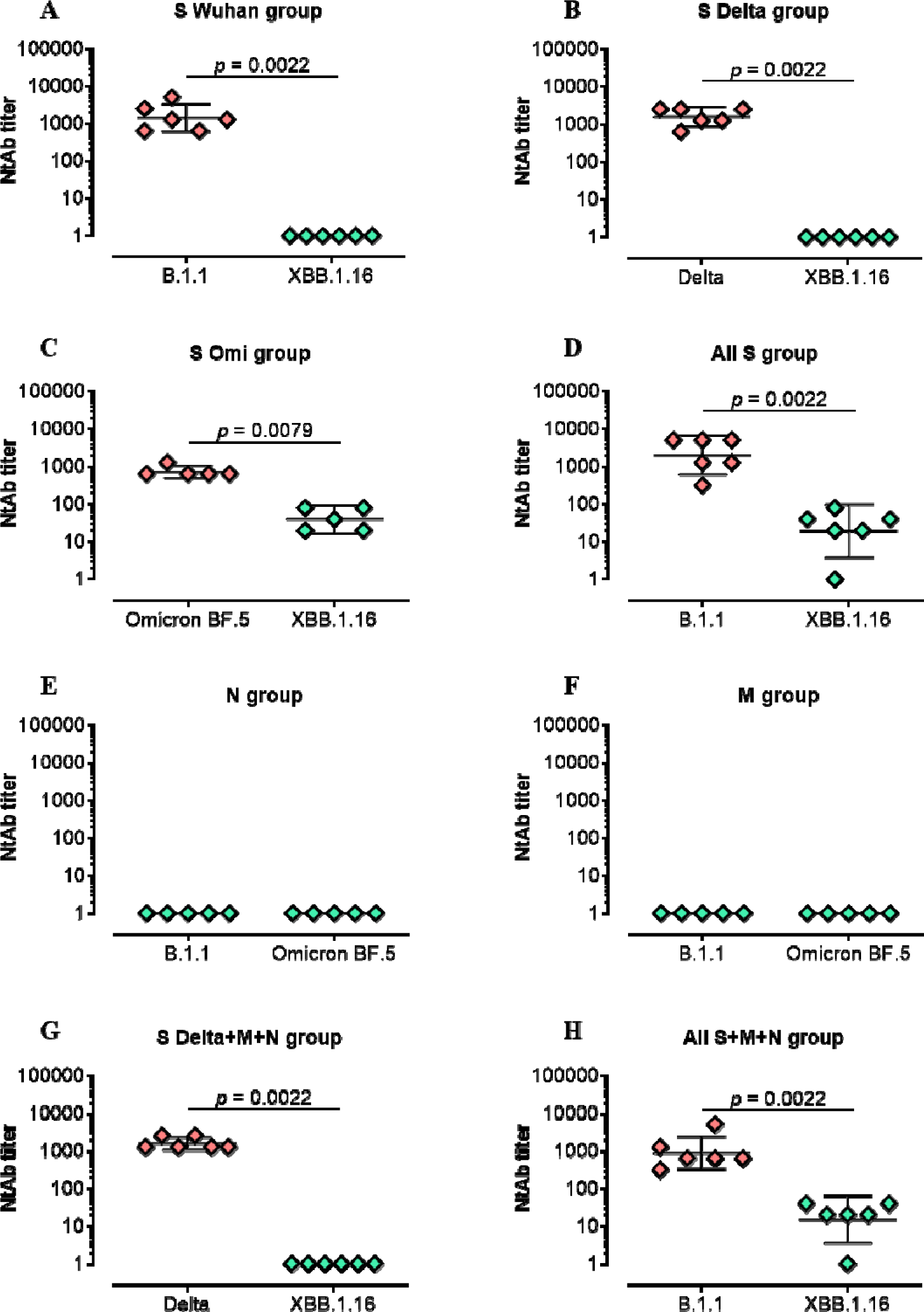
Serum neutralizing activity against B.1.1, Delta, Omicron BF.5, or XBB.1.16 in vaccinated BALB/c mice. Neutralizing antibody (NtAb) titers were determined by the highest pla ma dilution protecting 80% of the infected wells. Titer values are represented as scatter dot plots in logarithmic scale. Lines represent geometric means with 95% confidence interval. Mann-Whitney test was used for statistical analysis.

As expected, no neutralizing activity was detected in mice groups vaccinated with only mRNA-N or mRNA-M without addition mRNA-S in vaccine composition (**fig. 2E, F**).

The M-, N- and S-specific T cell responses were measured by intracellular cytokine staining (ICS) after stimulation of splenocytes with full lengths M, N proteins or RBD domain of Spike protein, respectively. Predominantly specific CD4+ T cell lymphocyte producing IL-2 and TNF-α were detected and significantly were higher compared to the PBS control group (**fig. 3**). Among mRNA-S vaccinated groups RBD-specific responses were observed in groups “All S” (*p* < 0.0001) and “S Delta+M+N” (*p* < 0.05) but not in “All S+M+N” group mostly when stimulated by RBD of Delta variant (**fig. 3A-F**). Similarly, N-specific CD4^+^ T cells predominantly expressed TNF-α and IL-2 compared to the PBS control group (*p* < 0.05) (**fig. 3G, H**). Among all mRNA-N vaccinated groups N-specific CD4^+^ T cell responses were more pronounced in “S Delta+M+N” group relative to the PBS control (*p* = 0.004 for TNF-α^+^CD4^+^ T cells, *p* = 0.0223 for IL-2^+^CD4^+^ T cells) (**fig. 3G, H**). For M protein, only IL-2 producing CD4^+^ T cells were in “S Delta+M+N” group compared with PBS control group (*p* = 0.0094) (**Fig. 3J)**.

**Figure 3.**
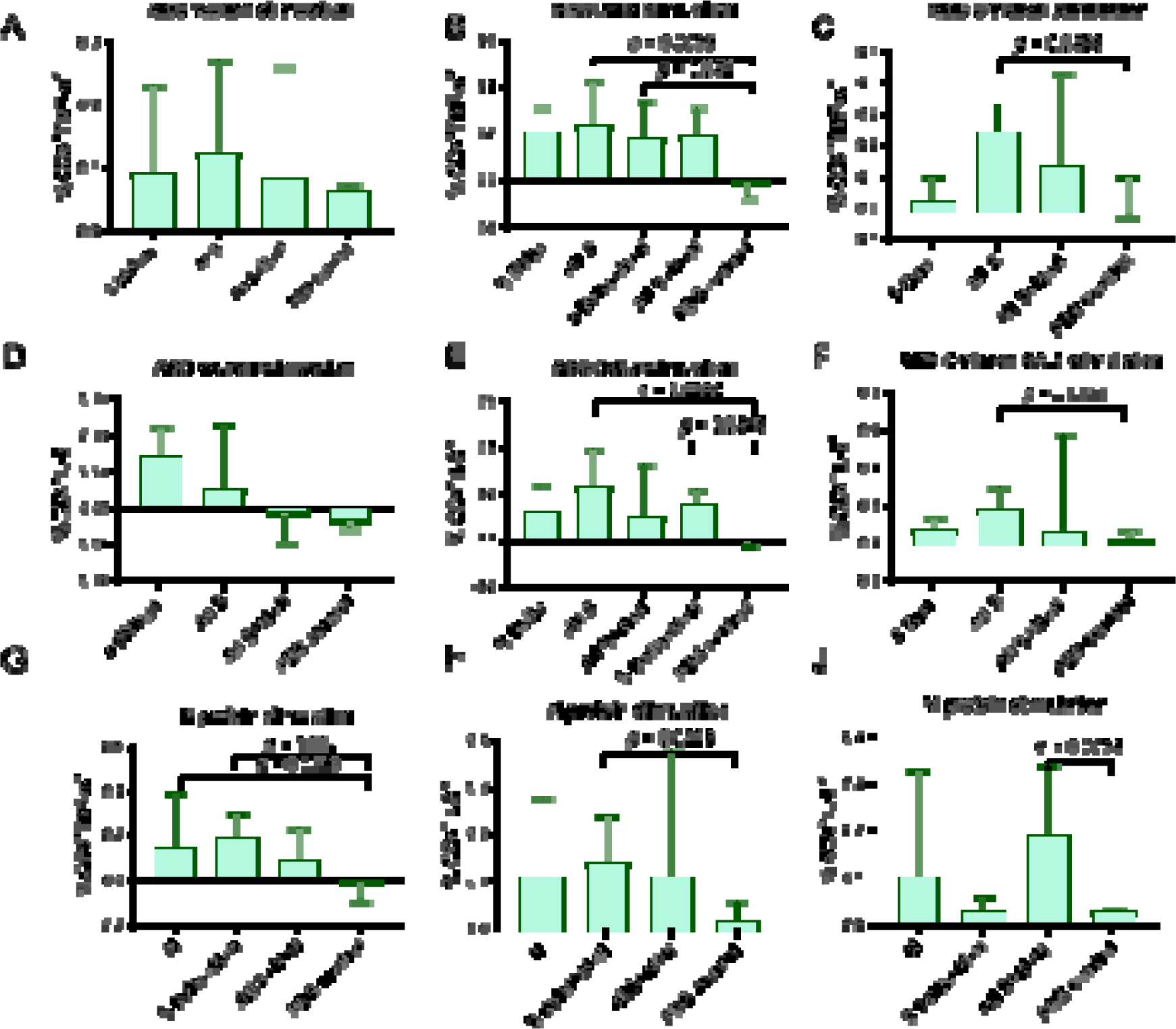
Vaccine-specific T cell responses in mouse spleen measured by ICS. Splenocytes were stimulated with full lengths N, M protein, or RBD domain of S protein, followed by immune staining and flow cytometry analysis. (**A-F**) Comparison of percentage of cytokine-positive, RBD-specific CD4^+^ T cells between vaccinated groups and PBS control. (percentage of total CD4^+^ T cells) (**G-J**) Comparison of percentage of cytokine-positive, N, M-specific CD4^+^ T cells between vaccinated groups and PBS control. Data are presented as median with 95% confidence interval. Kruskal-Wallis test and the post hoc Dunn’s multiple comparisons test was used for statistical analysis.

The data obtained from immunogenicity experiment have shown potential of including mRNA-N rather mRNA-M in mRNA-LNP vaccine. Furthermore, compared with mRNA-N alone, combination mRNAs coding N and S proteins in vaccine composition allows inducing more robust immune response to N protein from both humoral and cell-mediated immunity.

Then we evaluated protective efficacy of mRNA-LNP vaccines coding S and N proteins compared with commonly used and approved types of mRNA-LNP vaccines - coding S alone or two variants of S protein (bivalent mRNA-LNP vaccine). For s ch comparing, we used mRNA coding S protein from recently circulated Omicron XBB.1. For bivalent mRNA-LNP vaccine mRNA coding S protein of Wuhan strain was added to mRNA-S XBB.1. The mRNA-LNPs combinations for immunization and group names are listed in **figure 4A**. Vaccine efficacy was studied on transgenic mice, lineage B6.Cg-Tg(K18-AC 2), expressed ACE receptor for virus entry. Experimental design is represented in **figure 4B**. Three groups of mice were immunized at week 0 (prime) and week 3 (boost) were vaccinated ith mRNA-LNPs (5 μg of each mRNA-LNP per dose), phosphate-buffered saline was injected to the mice control group (PBS control). Then, mice were challenged intranasal with the SA S- CoV-2 Omicron BF.7 strain (10^5^ TCID_50_) at week 7. Viral titers and RNA copies in the lungs were measured 3 days after challenge (**Fig. 4B**). Compared to PBS control group, all mRNA-LNP compositions (mRNA-S alone, combined with mRNA-N or bivalent vaccine) induced total viral control with no detectable infectious virus in the lungs (*p* = 0.0007) ( **Fig. 4C**). In addition, all mRNA-LNP compositions demonstrated high efficacy in reducing copies of viral RNA in the lungs compared to the control group PBS (*p* < 0.05) (**Fig. 4D**). However, neither viral titers measurement nor quantification of viral RNA by the more-sensitive reverse transcription polymerase chain reaction (RT-PCR) approach did not reveal

**Figure 4.**
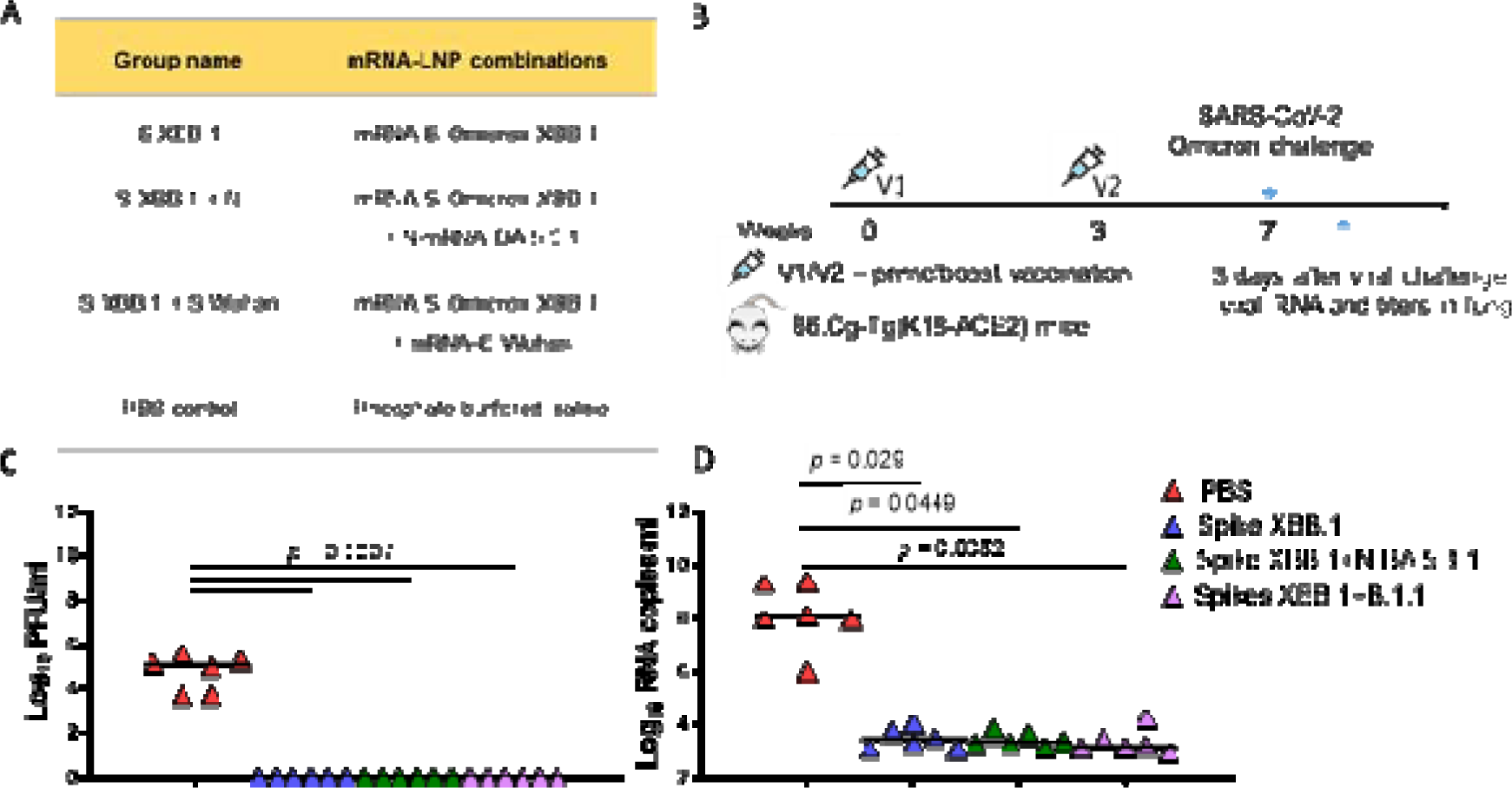
Mice SARS-CoV-2 challenge model. (**A**) List of vaccine-candidates compositions and group names. (**B**) Experimental design of challenge experiment. Four groups of B6.Cg-Tg(K18-ACE2) mice (*n* = 15 per group) were intramuscularly immunized with mRNA-LNP combinatio (5μg of each mRNA per dose) or PBS (control group) at weeks 0 and 3 and were intranasally challenged with an SARS-CoV-2 BF.7 strain (10^5^ TCID_50_) at week 7. On 3 day after challenge (*n* = 6) lung tissues were harvested for analysis of viral RNA copies, viral titers. (**C**) Comparison of viral titers (plaque-forming unit (PFU)/ml) in the mouse lungs between different groups. (**D**) Comparison of viral RNA copies in the mouse lungs (log_10_ RNA copies per/ml) between different mice groups. Data are represented as scatter dot plots in logarithmic scale. Lines represents medians, Kruskal-Wallis test and the post hoc Dunn’s multiple comparisons test was used.

## DISCUSSION

Recently, we have developed an mRNA-LNP platform providing efficient long-term expression of the encoded gene *in vivo* through both intramuscular and intravenous administration. In this study we aim to demonstrate the potential of our mRNA-LNP platform for the development of COVID-19 vaccine candidates.

Based on our platform, we obtained mRNA-LNP variants that encode the structural proteins of SARS-CoV-2. We conducted comprehensive investigations into the immunogenicity and protective efficacy of mRNA-LNP encoding M, N, or S proteins, both individually and in various combinations. We also examined how they mutually influence each other in eliciting a specific immune responses.

It is known that mRNA vaccines against COVID-19 are capable of inducing robust CD4+ and CD8+ T cell responses and strong antibody responses [24][25]. The data obtained in our study demonstrated high potential of using our mRNA-LNP platform in vaccine development. All studied vaccine compositions encoded S and N proteins induced high GMT of RBD- and N-specific IgG antibody in mice (GMT for anti-RBD IgG was averaged 10^6^). In addition, T cell responses mainly as specific CD4+ T cell lymphocyte producing IL-2 and TNF-α were detected in response to RBD or full-length N protein stimulation. Contrary to expectations, poor immune response was observed when mice were vaccinated with mRNA-LNP coding M protein. However, limited data confirms its immunogenic properties caused humoral [26] and strong T cell [27][15] responses.

We also obtained quite expected results of virus-neutralizing activity in vaccinated mice. High neutralizing activity was detected in groups vaccinated with alone mRNA-S or combined of three mRNA-S against closely related strains but was not detectable or significantly lower against evolutionarily distant variant Omicron XBB.1.16. The data reflects the problem of the ongoing evolution of SARS-CoV-2 in the human population resulted in the emergence of variants of concern (VOCs). Until late 2021, before the emergence of Omicron, VOCs such as Alpha and Delta were mainly associated with increased transmissibility and modest degrees of immune escape. However, Omicron had more than a few dozen extra mutations in the Spike gene (especially in the RBD) and that gave the advantage in immune escape properties which in turn led to displacement of Delta by Omicron. Further substitutions that arose in these Omicron sub-lineages (BA.2, BA.4, BA.5, XBB), also mainly clustering in the RBD, have caused significant reductions in the antibody neutralization titers of sera from individuals who are naturally infected or vaccinated [28].

Today, due to the ongoing emergence of new VOCs, the global vaccine strategy aimed at updating the S-protein in vaccines (the introduction of bivalent or monovalent boosters) still remains the most relevant. In addition, in the face of constantly changing S protein antigenic properties, design and selection of VOC-specific sequences are too slow to be ideal. However, the development of next-generation vaccine strategies for broader and long-lasting protection is equally important.

For this purpose several studies have tested SARS-CoV-2 candidate vaccines that include other structural proteins (mostly the N protein) in adenoviral vectors [19][29], modified vaccinia Ankara vectors [30][31], subunit protein vaccine candidate [32][33] and others [34]. These studies showed protective efficacy, T-cell and humoral response caused by vaccination. However, they did not compare efficacy with the current, clinically proven S-targeting vaccines. In addition, most of these vaccine approaches were only tested against earlier circulating VOCs, and their efficacy against the predominant Omicron remains unclear.

As for mRNA vaccines in one study the authors created mRNA-LNP that encodes the full-length N and S proteins of SARS-CoV-2 (Wuhan-Hu-1 strain). They showed that compared to mRNA-S alone, combination mRNA-S+N vaccination induced more robust control of the Delta and Omicron variants in the lungs and also provided enhanced protection in the upper respiratory tract [35].

In addition, some interesting results were obtained in humoral and cell responses in our study. Although, mRNA-S and mRNA-N doses in all used vaccines were identical (5 μg) some differences were found in the strength of the immune response to expressed antigens between groups. Our data suggest that the addition mRNAs encoding S and M antigens to the mRNA-N in the vaccine composition enhanced immunogenicity of mRNA-N inducing more robust humoral response to the N protein. However, the addition mRNAs encoding S and N antigens to the mRNA-M did not affect production of M-specific binding antibodies. Furthermore, the addition of mRNAs encoding N and M proteins to mRNA-S vaccine preparation did not affect humoral response caused by mRNA-S.

Similar pattern revealed in T cell responses after stimulation of splenocytes with full lengths N protein. Among all mRNA-N vaccinated groups N-specific CD4^+^ T cell responses were more pronounced in “S Delta+M+N” compared with group vaccinated mRNA-N alone. Based on our binding antibody assay results, we hypothesize that in our compositions the S protein plays a key role in enhancing the anti-N protein immune response.

The same phenomena but with the opposite effect was observed earlier in another study by Hajnik et al. [35]. The authors generated mRNA-LNP vaccines that encodes the full-length N and S proteins of SARS-CoV-2 (Wuhan-Hu-1 strain). They also showed that mRNA-N alone in vaccine was highly immunogenic and induced robust N-specific T cell responses and binding antibodies. In contrast to our results, vaccination with the combination of mRNA-S+N led to augmented S-specific immunity in T-cell response compared to mRNA-S alone. However, the authors did not compare mRNA-N to mRNA-S+N vaccine combination in the same experiment as we did. Like us, the authors find it difficult to explain this phenomenon. One hypothesis is that cross-priming effects occur between N and S antigens after vaccination or that mRNA-N co-immunization may induce an immune environment that favors the generation of S-specific immunity (or, conversely, as we obtained in our study).

In another study [36] it was shown the additional enhancement of T-cell response to the structural M and N SARS-CoV-2 proteins after mRNA vaccine in individuals with past SARS-CoV-2 infection and infection-naïve individuals with cross-reactivity to the seasonal coronaviruses. Unexpectedly, the cellular response was not exclusively directed toward S-related epitopes after spike mRNA vaccination. It is still controversial how exclusively specific T-cell antigen recognition is [37], and reports are suggesting that T-cell recognition might be highly promiscuous with individual T-cell clones being able to cross-reactively recognize different epitopes [38].

Thus, more detailed studies of mechanisms of S-, N-, M-specific immunity formation after combined mRNAs vaccination is needed, such as protein expression, antigen presentation, and stimulation of the innate response, formation of pool of T- and B cells, their clone diversity and interaction. Understanding the development of adaptive immunity to the SARS-CoV-2 virus is essential for vaccine development and setting respective pandemic control measures in light of the observed constant evolution of the pathogen.

In the challenge experiment we compared mRNA-LNP vaccine coding S and N proteins with commonly used and approved types of mRNA-LNP vaccines - coding S alone or bivalent mRNA-LNP vaccine. All mRNA-LNP compositions have shown excellent results in protective efficacy: total viral control with no detectable infectious virus in the lungs and reducing copies of viral RNA in the lungs compared to the control group PBS three days after intranasal challenge with Omicron BF.7 strain. mRNA-LNP vaccine coding alone S protein from XBB.1 has sufficient protective properties without adding mRNA-S coding ancestral strain (bivalent vaccine) or N protein. However, we did not investigate the protective efficacy these same vaccine compositions when mice were challenged with evolutionarily distant variant (for example, Delta, or new highly mutated BA.2.86). That is our main limitation. Conducting a similar experiment in the future could help to see the difference between compositions and possible benefits of adding the more conservative N protein in our mRNA-LNP vaccine.

## CONCLUSIONS

Our study serves as a compelling demonstration of the potential utility of our mRNA-LNP platform in vaccine development, using SARS-CoV-2 infection as an illustrative example. The mRNA-LNP vaccine compositions encoding the structural proteins (S and N) have demonstrated robust immunogenicity in mice, eliciting both humoral and cellular immune responses. Additionally, we have elucidated that the presence of different protein-encoding mRNAs within the mRNA-LNP vaccine compositions could exert mutual influence, resulting in variable degrees of immune response strength compared to mRNA encoding individual proteins.

In addition, our mRNA vaccine-candidates encoding the S protein demonstrated high virus neutralizing activity against homologous variants. In the case of the Omicron XBB.1 S- antigen, protective efficacy is provided not only against the early Omicron BF.7 variant, but also against the currently dominant XBB.1.16. Expression of this antigen, alone or in combination with other antigens, effectively provides protection against the Omicron BF.7 strain. Updating S- glycoprotein vaccine formulations to account for circulating variants of SARS-CoV-2 remains the main strategy for obtaining effective vaccine formulations until a new strategy is developed that can generate a broadly neutralizing immune response.

## MATERIALS AND METHODS

### mRNA synthesis and LNP formulation

mRNAs encoding following antigens were synthesized:

- S protein of wild-type strain SARS-CoV-2 (Wuhan)
- S protein of Delta variant
- S, N and M proteins of Omicron variant (BA.5.3.1)
- S protein of Omicron variant (XBB.1)

mRNA-LNP preparations were produced as previously described [22]. Briefly, the coding regions for investigated antigens (M, N, S proteins) were cloned into a vector for *in vitro* transcription (IVT) based on the pJAZZ-OK linear bacterial plasmid. Cloning and plasmid production were performed using *E. coli* BigEasy™-TSA™ Electrocompetent Cells (Lucigen). All cloning procedures were verified by Sanger sequencing using BigDye® Terminator v3.1 Cycle Sequencing kit on 3500 Genetic Analyzer (Applied Biosystems). The pDNA for IVT were isolated, purified from a culture of the *E.coli* and then digested with BsmBI-v2.

IVT was performed as described earlier [22]: 100-μl reaction volume contained 3 μg of DNA template, 3 μl T7 RNA polymerase (Biolabmix) and 10xBuffer (TriLink), 4 mM trinucleotide cap 1 analog ((3′-OMe-m7G)-5′-ppp-5′-(2′-OMeA)pG) (Biolabmix), 5 mM m^1^ΨTP (Biolabmix) replacing UTP, and 5 mM GTP, ATP and CTP. After 2 h incubation at 37 °C, 6 μl DNase I (Thermo Fisher Scientifiс) was added for additional 15 min, followed by mRNA precipitation with 2M LiCl (incubation for 1 h in ice and centrifugation for 10 min at 14,000 g, 4°C) and carefully washed with 80% ethanol. RNA integrity was assessed by electrophoresis in 8% denaturing PAGE.

*In vitro* transcribed mRNAs were encapsulated in LNPs as described earlier with some modifications (CC). Briefly, lipids were dissolved in ethanol at molar ratios of 46.3:9:42.7:1.6 (ionizable lipid (ALC-0315): distearoyl PC:cholesterol: PEG-lipid (ALC-0159). The lipid mixture was combined with 10 mM sodium citrate buffer (pH 3.0) containing mRNA (0.2 mg/ml) at a volume ratio of 3:1 (aqueous phase:organic phase) using a microfluidic cartridge technology in NanoAssemblr Ignite device (Precision NanoSystems). Formulations were then dialyzed against PBS buffer (VWR Life Science) through a 3.5 K MWCO Slide-A-Lyzer dialysis cassette (Thermo Scientific), followed by passed through a 0.2-μm filter and stored at 4 °C until use.

The diameter and size distribution of the mRNA-LNP was measured by dynamic light scattering (DLS) using a Zetasizer Nano ZS instrument (Malvern Panalytical). The mRNA-LNP suspension diluted in water 20 times was loaded into a cuvette for a measurement at 25 °C. LNPs in PBS were read using the solvent parameter ‘water’. One measurement consists of three readings and each reading derives from 13 acquisitions. The DLS data presented for each mRNA-LNP preparation is the average value of these four readings.

The mRNA encapsulation efficiency and concentration were determined by SYBR Green dye (SYBR Green for PCR, Lumiprobe) followed by fluorescence measurement. Briefly, mRNA-LNP samples were diluted with TE buffer (pH8.0) in the absence or presence of 2% Triton-X-100 in a black 96-well plate. Standard mRNA (4 ng/μL) was serially diluted with TE buffer in the absence or presence of 2% Triton-X-100 to generate standard curves. Then the plate was incubated 10 min followed by addition of SYBR Green dye (100 times diluted) to each well to bind RNA. Fluorescence was measured by at 454 nm excitation and 524 nm emission using Varioscan LUX (Thermo Fisher Scientifiс). The concentrations of mRNA after LNP disruption by Triton-X-100 (C _total mRNA_) and before LNP disruption (C _outside mRNA_) were determined using corresponding standard curves. The concentration of mRNA loaded into the LNP was determined as the difference between the two concentrations multiplied by the dilution factor of the original sample. Encapsulation efficiency was calculated by the formula: (E%)L=L(C _total_ mRNA)–(C outside mRNA)]/ (C total mRNA) ×L100%.

### Cell transfection

The expression of each vaccine-encoded viral protein from the *in vitro* transcribed mRNA was confirmed in HEK293T cells (ATCC). Cell transfection was performed as previously described [22]. Briefly, 3×10^4^ cells per well were transferred into the 96-well plates (Greiner) in 75 μl of DMEM (Paneco) supplemented with 10% FBS (HyClone) in the presence of penicillin and streptomycin (Paneco). Cell culture was incubated at 37 °C in 5% CO_2_ atmosphere. Next day, cell culture was transfected with the mRNA. For this, 30 ng of mRNA in 20 μl Opti-MEM (Gibco) per well was mixed with a solution of 0.06 μl of GenJector^TM^-U (Molecta) in 3 μl Opti-MEM (per well), incubated for 15 min, added to the cells. Transfected cells were incubated at 37 °C with 5% CO_2_ for 24 h. Then cells were collected, centrifuged at 300 g, 5 min, +4 °C. Supernatant was removed following the addition of 100 µl of NENT lysis buffer to the cell pellet and incubation on ice for 30 min. Cell lysate was centrifuged at 13000g 5 min, +4 °C, and supernatant were subsequently analyzed.

### Detection of protein production

Protein production was confirmed by in-house bead-based immunoassay (Luminex® xMAP® technology). MagPlexTM-C carboxylated microspheres (beads) were used for conjugation. Mouse monoclonal N- and S- specific IgG antibodies (Hytest, clone RBD5308 and clone C518) and rabbit polyclonal M-specific IgG (Antigenic, #P0DTC5) were conjugated to beads (each to his own beads region) according to the protocol for two-step carbodiimide reaction in the Luminex Cookbook [23]. Briefly, the coupling procedure was performed as follows: 1 ×10^6^ beads of one region were activated with 10 µL of 50 mg/mL EDC (Thermo Fisher Scientific) and 10 µL of 50 mg/mL s-NHS (ThermoFisher Scientific) in 80 µL activation buffer (0.1 M NaH_2_PO_4_, pH 6.2) for 20 min at 25 °C. After that, the activated beads were washed twice and resuspended in 500 µL of coupling buLer (50 mM MES, pH 5.0) with the 10 µg of each antibody. The incubation was conducted for 2 h at RT in the dark and mixed on a Rotamix RM-1L rotator (ELMI, Riga, Latvia). After the coupling procedure, the beads were washed three times, resuspended in 1 mL PBS-TBN buffer (PBS, 1% BSA, 0.1% Tween-20, 0.05% NaN_3_, pH 7.4), quantified using an automatic cell counter TC-20 (Bio-RAD, Hercules, US) and stored in the dark at 2–8 °C until use.

The immunoassay was performed in monoplex as previously described with some modifications [39]. For reference standards recombinant proteins N, M or RBD domain were used (see below). The reference standards was 3-fold serially diluted over 8 points using PBS-TBN buffer for establishing the calibration curve. Standard dilutions and cell lysate were 20-fold diluted with PVXC buffer (PBS, 0.8% PVP, 0.1 % casein, 0.5% *PVA,* 0.05% NaN_3_). Then, 2500 microspheres in 80 µL PBS-TBN buffer per well were incubated with 20 µL of sample (final dilution of 1:100 for all samples in the well) for 60 min at 37 °C in the dark on a rotating shaker (800 rpm). Then, beads were washed twice with 200 µL PBS-TBN buffer using an Agilent BioTek 405 TS Microplate Washer magnetic plate separator (Agilent Technologies, USA). Next, 100 µL of biotinylated secondary antibodies to corresponding proteins in PBS-TBN buffer (4 µg/mL) was added to microspheres and incubated for 60 min in the same conditions. Following secondary antibodies were used: monoclonal N- and S- specific IgG antibodies (Hytest, rabbit IgG, clone C706 and mouse IgG, clone RBD5305, respectively), and rabbit polyclonal anti-M- protein antibodies (Atagenix, #PODTC5). Then, beads were washed twice in the same manner. Next, 100 µL of SAPE (Thermo Fisher Scientific) in PBS-TBN buffer (4 µg/mL) was added to microspheres and incubated for 30 min in the same conditions. After a final wash step, beads were resuspended in 100 µl of PBS-TBN and analyzed on a MAGPIX instrument. For each identified analyte, the MFI-value was converted to ng/mL by interpolation from a 5-parameter logistic (5-PL) curve of reference standard using the MILLIPLEX® Analyst 5.1 software (The Life Science/Merck KGaA).

### Mouse immunization and SARS-CoV-2 challenge

All animal experiments were performed in accordance to the Directive 2010/63/EU (Directive 2010/63/EU of the European Parliament and of the Council of 22 September 2010 on the Protection of Animals Used for Scientific Purposes. Off J Eur Communities L276:33–79.), FELASA recommendations [40] and the Inter-State Standard of «GLP» (GOST - 33044-2014) approved by the Institutional Animal Care and Use Committee (IACUC) of the Federal Research Centre of Epidemiology and Microbiology named after Honorary Academician N.F. Gamaleya and were performed under Protocol #56 from 31 July 2023. All persons using or caring for animals in research underwent training annually as required by the IACUC.

Vaccine immunogenicity was evaluated in 4-5-week-old females BALB/c mice (The Federal Medical-Biological Agency or FMBA, Stolbovaya breeding nursery, Russia). For immunogenicity, nine groups of mice (six per group) were immunized intramuscularly at week 0 (prime) and week 3 (boost), respectively. Mice groups were immunized with either PBS (control group), mRNA-S-Wuhan (5 μg), mRNA-S-Delta (5 μg), mRNA-S-Omicron (5 μg), mRNA-N (5 μg), mRNA-M (5 μg), or combined mRNA vaccines (5 μg for each mRNA component) – mRNAs S-Delta+M+N, mRNAs S-Wuhan+S-Delta+S-Omicron (mRNA-all-S group), or mRNAs S-Wuhan+S-Delta+S-Omicron+M+N (mRNA-all-antigens group), at week 0 (prime) and week 3 (boost), respectively. The vaccine or PBS was administered at 100 μl per injection. Blood and serum samples were collected 2 weeks after prime vaccination (V1) and 2 weeks after booster vaccination (V2) to measure vaccine-induced binding antibody response. All mice were euthanized 2 months after booster vaccination (V2). Blood and serum and spleen samples were collected for analyses of vaccine-induced humoral (neutralizing antibodies) and cellular immune responses.

Vaccine efficacy was studied in SARS-CoV-2 challenge on 6-week-old transgenic B6.Cg-Tg(K18-ACE2) mice (“Medgamal”subsidiary of N. F. Gamaleya Federal Research Center for Epidemiology and Microbiology). Mice were divided into five groups of 16 individuals per group (8 males and 8 females). Mice groups were immunized with either PBS (placebo group), mRNA-S XBB.1 (5 μg), or combined mRNA vaccines (5 μg for each mRNA component) – mRNAs-S XBB.1+N, or mRNAs S-Wuhan+S-XBB.1, at week 0 (prime) and week 3 (boost), respectively. The vaccine or PBS was administered at 100 μl per injection. Two weeks after booster vaccination, all mice were transferred to animal biosafety level 2 facility. Then, after two weeks, mice were challenged intranasally with an SARS-CoV-2 BF.7 strain (10^5^ TCID_50_). Four days after viral challenge, six mice from each group were euthanized and equivalent portions of the lung tissues were collected for quantification of SARS-CoV-2 viral loads. Mice body weights and survival were monitored daily to evaluate vaccine-induced protection for two weeks after challenge.

### Binding IgG detection by ELISA

Vaccine-induced, binding IgG against the N, M proteins and receptor-binding domain (RBD) of S protein were measured by “in-house” ELISA as previously described [41]. Briefly, plates (Xema, Moscow, Russia) were coated with recombinant RBD of different SARS-CoV-2 variants (0.5 μg/ml; EVV00312 (Delta) and EVV00329 (BA.5), AntibodySystem; 8COV1 (Wuhan), HyTest, Russia), N protein (1 μg/ml “In-house” production) or M protein (1 μg/ml; ATEP02465COV, Atagenix Laboratories). The next day, the plates were blocked for 2 h at room temperature with a S002X buffer (Xema, Moscow, Russia). After blocking buffer was removed, serially diluted serum samples were added into the wells and incubated for 1 h at 37 °C (initial dilution, 1:100; 1:2 serial dilution in ELISA buffer S011 (Xema, Moscow, Russia). Plates were washed 3 times and incubated with 100 μL HRP-conjugated anti-mouse IgG secondary antibody (L20/01; HyTest; 1:25000) for 1 hour at 37°C. After final wash, plates were visualized using chromogen-substrate solution R055 (Xema, Russia). After 10 min, the reaction was stopped using a 10% HCl. Plates were read at 450 nm wavelength within 15 min by using Multiscan reader (Thermo Scientific, MA, USA). Binding IgG endpoint titers (EPTs) for each sample were calculated.

### ICS and flow cytometry

Isolated mouse splenocytes were washed with fluorescence-activated cell sorting (FACS) buffer (PBS, 0.5% BSA) and resuspended in RPMI 1640 supplemented with 10% FBS, 2-mercaptoethanol, penicillin-streptomycin, and L-glutamine. 2 ×10^6^ cells/ml were stimulated with recombinant full-length proteins (10 μg/ml; in-house production N protein or M protein, ATEP02465COV, Atagenix Laboratories) or RBD domain (10 μg/ml) of different SARS-CoV-2 variants (8COV1 (Wuhan strain), HyTest, Russia; EVV00312 (Delta) and EVV00329 (BA.5), AntibodySystem). Stimulation was performed in the presence of anti-CD28 (0.5 μg/ml; Invitrogen eBioscience™ Anti-Mouse CD28 Syrian hamster / IgG 37.51) and anti-49 (0.5 μg/ml; Invitrogen eBioscience™ CD49d (Integrin alpha 4) Monoclonal Antibody, Functional Grade, Rat / IgG2b, kappa (R1-2)) and Brefeldin A (5 μg/ml; Sigma-Aldrich) for 16 hours. Cells stimulated with phorbol 12-myristate 13-acetate and ionomycin were included as positive control. After stimulation, cells were washed, then anti CD16 /32 were added for Fc- receptors blocking on ice for 10 minutes (Biolegend, Purified anti-mouse CD16/32 AntibodyRat IgG2a, λ 93). Then cells were first stained stained with LIVE/DEAD aqua viability dye (Life Technologies, Carlsbad, CA, USA), anti-mouse fluorochrome-conjugated antibodies to surface markers, including anti-CD3-BV421 (Biolegend, Brilliant Violet 421™ anti-mouse CD3 Antibody, Rat IgG2b, κ, Clone 17A2), anti-CD8-BV650 (Biolegend, Brilliant Violet 650™ anti-mouse CD8a Antibody, Rat IgG2a, κ, 53-6.7), anti-CD4- BV785 (Biolegend, Brilliant Violet 785™ anti-mouse CD4 Antibody, Rat IgG2a, κ, RM4-5) for 20 minutes in the dark at room temperature. After wash step cells were fixed and permeabilized using IntraPrep Permeabilization Reagent (Beckman Coulter, Brea, CA, USA). Subsequently, cells were washed again and stained with anti-mouse antibodies to intracellular markers anti-TNF-α-Alexa Flour 700 (Biolegend, Alexa Fluor® 700 anti-mouse TNF-α Antibody, Rat IgG1, κ, MP6-XT22), anti-IFN-γ-Alexa Flour 480 (Invitrogen Alexa Fluor™ 488, eBioscience™ anti-mouse IFN gamma Antibody Rat / IgG1, kappa XMG1.2) and anti-IL-2-PE (Affymetrix eBioscience, PE anti-mouse IL-2 Antibody, IgG2b κ, JES6-5H4) for 20 minutes in the dark at room temperature. Cells were then washed and were analyzed by a Cytoflex S flow cytometer (Beckman Coulter, Brea, CA, USA). Analysis of the flow cytometry data was performed using CytExpert software version 2.3.

### RNA extraction and RT-PCR quantification of viral RNA copies

Lungs were harvested from mice 3 days post-infection. Following harvest, lungs were weighed, and then homogenized in sterile DMEM with gentamycin to generate a 20% lung-in-medium solution. Total RNA was extracted from lung homogenates using the ExtractRNA Reagent (Eurogen, Moscow, Russia) following the manufacturer’s instructions. Amplification and quantification of influenza A virus RNA were carried out by using a one-step RT-qPCR technique.

### Virus-neutralization assay

Vero E6 cells were plated in 96-well plates at a density of 3 × 10^4^ cells per well. After 18 h incubation, different dilutions of the compound in DMEM with 2% FBS were added to the cell monolayer in triplicate and incubated for 1 h at 37 °C. Then, the cells were infected with the corresponding SARS-CoV-2 virus strain at 100 TCID50. The virus-induced CPE was evaluated after 72–96 h of infection using an MTT method. The data obtained from the experiment were analyzed using GraphPad Prism 8.0 software.

### Statistical analysis

Statistical analysis was performed using the GraphPad Prism 8.0 software. Nonparametric tests were used throughout this paper for statistical analysis. Data were presented as geometric mean with 95% confidence interval or as median. Comparison among groups was performed by using either Mann-Whitney test (two groups) or Kruskal-Wallis test (multiple comparison) with Dunn’s post hoc test. Two-tailed *P* values were denoted, and *p*<0.05 values were considered as significant.

## Acknowledgments

We are grateful to Timofey A. Remizov and Anastasia A. Zakharova for their help with the purchase of reagents and paperwork for ongoing projects. We are grateful to Igor V. Grigoriev for assistance with ELISA and Maria S. Poponova for cell assay implementation.

## SUPPLEMENTARY

**Figure S1.**
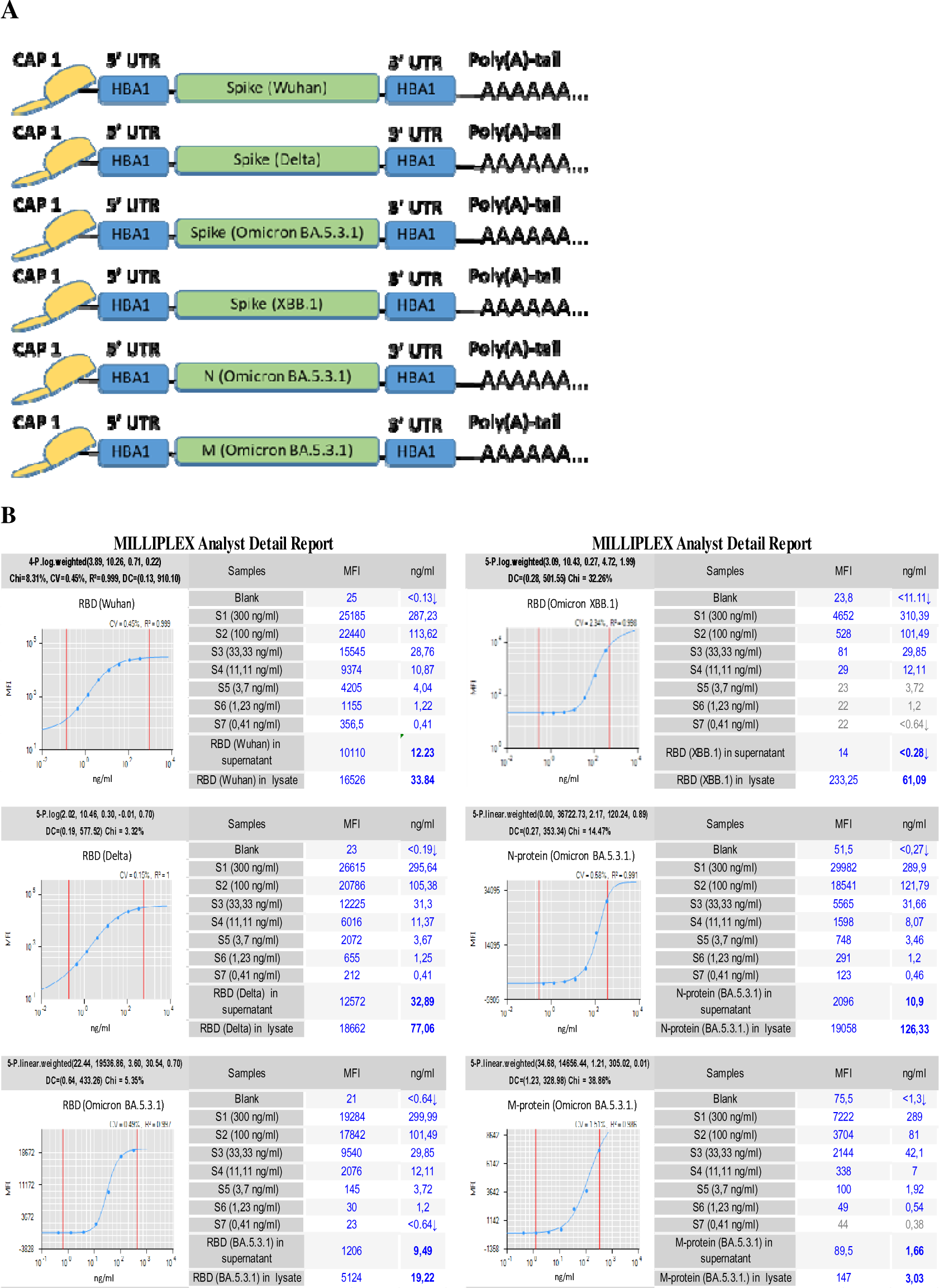
Schematic representation of mRNAs, encoding SARS-CoV-2 structural proteins using in this study (**A**) and the check of mRNAs translation on HEK293T (**B**). Bead based immunoassay results. Median fluorescent intensity (MFI) was measured on a MAGPIX instrument and converted in ng/ml. For each identified analyte (RBD, N or M), the MFI-value was converted to ng/mL by interpolation from a 5-parameter logistic (5-PL) curve of reference standard using MILLIPLEX® Analyst 5.1 software (The Life Science/Merck KGaA).

**Table S1.**
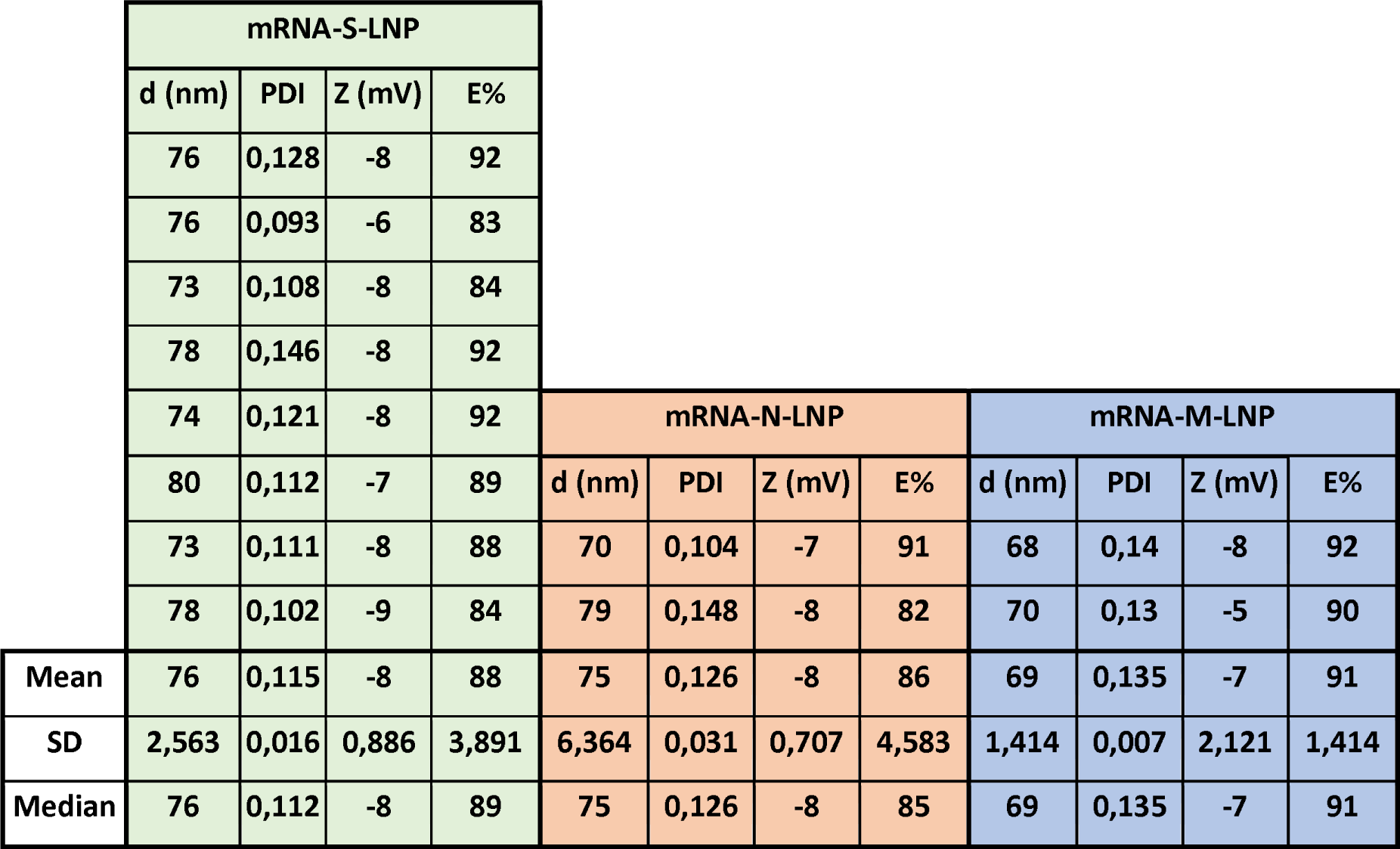
Physicochemical properties of mRNA-LNP formulations.

**Figure S2.**
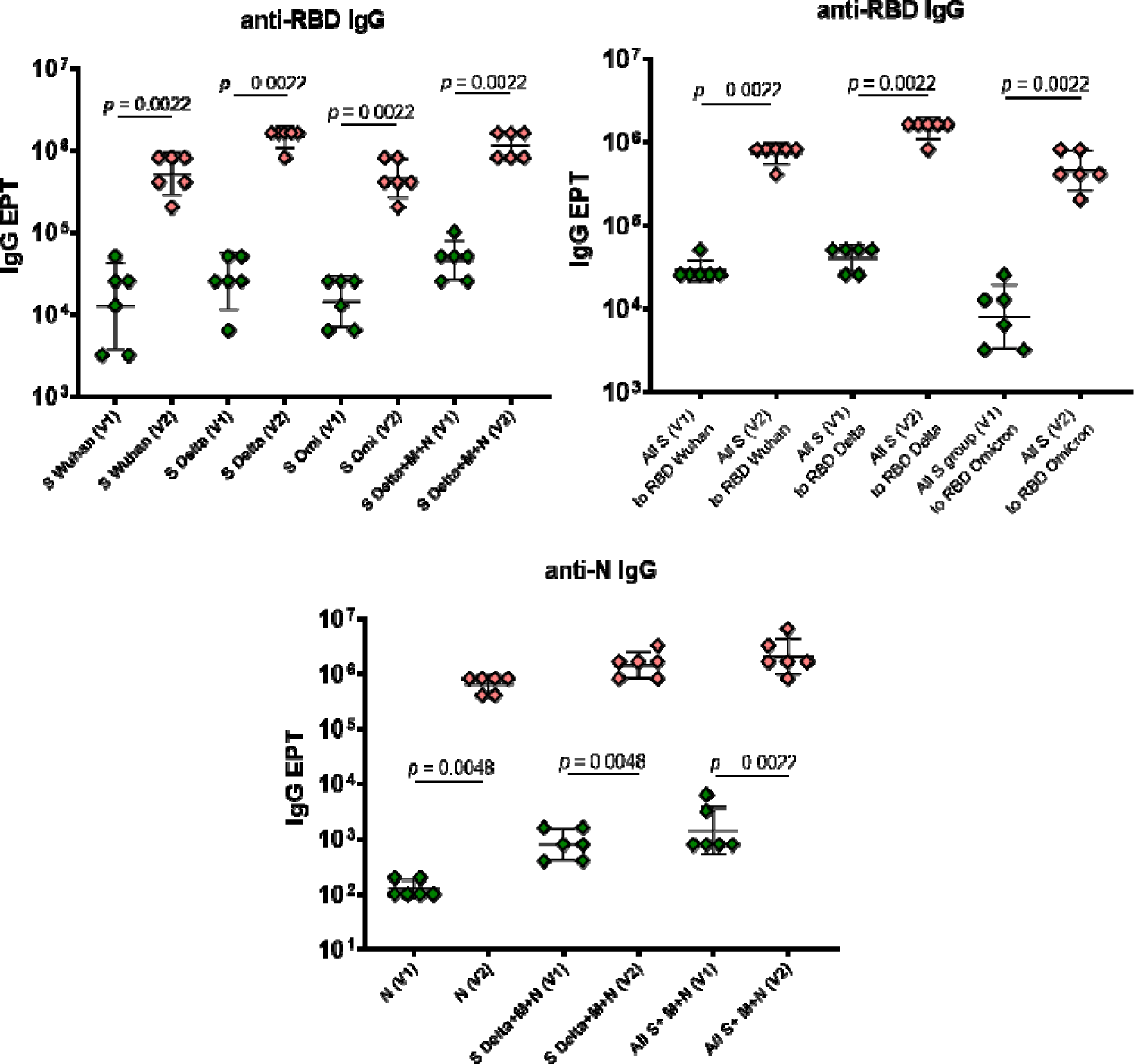
ELISA results are shown for serum N-, RBD-specific binding IgG after prime (V1) and boost (V2) vaccinations. EPT values are represented as scatter dot plots in logarithmic scale. Lines represent geometric means with 95% confidence interval. Mann-Whitney test was used for statistical analysis.

**Figure S3.**
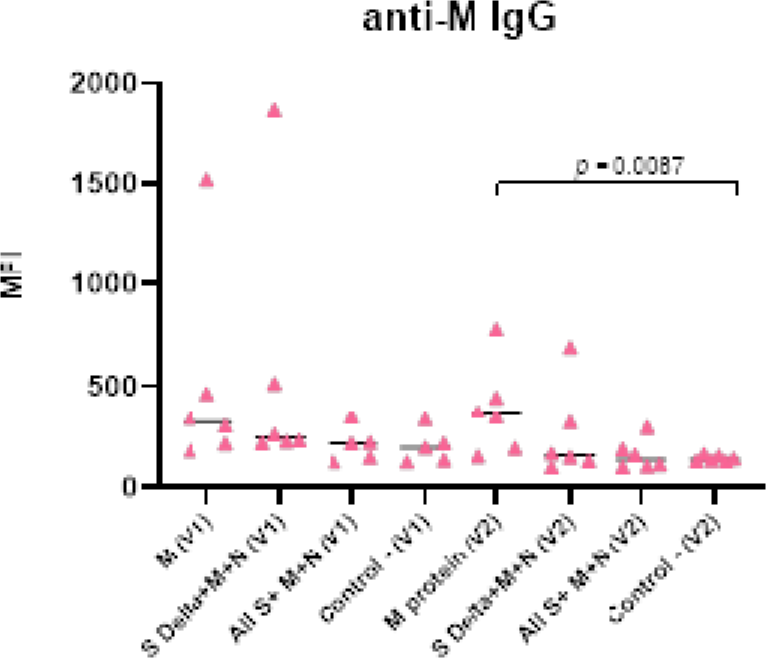
Bead based immunoassay results are shown for serum M-specific binding IgG after prime (V1) and boost (V2) vaccinations. MFI – median fluorescent intensity. Mann-Whitney test was used for statistical analysis.

